# *Chlamydomonas reinhardtii* Tubulin-Gene Disruptants for Efficient Isolation of Strains Bearing Novel Tubulin Mutations

**DOI:** 10.1101/2020.04.07.031005

**Authors:** Takako Kato-Minoura, Yutaro Ogiwara, Takashi Yamano, Hideya Fukuzawa, Ritsu Kamiya

**Affiliations:** Department of Biological Sciences, Faculty of Science and Engineering, Chuo University, 1-13-27 Kasuga, Bunkyo-ku, Tokyo 112-8551, Japan; Biological Science Course, Graduate School of Science and Engineering, Chuo University, 1-13-27 Kasuga, Bunkyo-ku, Tokyo 112-8551, Japan; Graduate School of Biostudies, Kyoto University, Kyoto 606-8502, Japan

**Keywords:** microtubule, herbicide, anti-cancer drug

## Abstract

The single-cell green alga *Chlamydomonas reinhardtii* possesses two α-tubulin genes (*tua1* and *tua2*) and two β-tubulin genes (*tub1* and *tub2*), with the two genes in each pair encoding identical amino acid sequences. Here, we used an *aphVIII* gene cassette insertional library to establish eight disruptants with defective *tua2, tub1*, or *tub2* expression. None of the disruptants exhibited apparent defects in cell growth, flagellar length, or flagellar regeneration after amputation. Because few tubulin mutants of *C. reinhardtii* have been reported to date, we then used our disruptants, together with a *tua1* disruptant obtained from the *Chlamydomonas* Library Project (CLiP), to isolate novel tubulin-mutants resistant to the anti-tubulin agents propyzamide and oryzalin. As a result of several trials, we obtained 8 strains bearing 7 different α-tubulin mutations and 24 strains bearing 12 different β-tubulin mutations. Some of these mutations are known to confer drug resistance in human cancer cells. Thus, single-tubulin-gene disruptants are an efficient means of isolating novel *C. reinhardtii* tubulin mutants.

**IMPORTANCE:** *Chlamydomonas reinhardtii* is a useful organism for the study of tubulin function; however, only five kinds of tubulin mutations have been reported to date. This scarcity is partly due to *C. reinhardtii* possessing two tubulin genes each for α- and β-tubulin. Here, we obtained several strains in which one of the α- or β-tubulin genes was disrupted, and then used those disruptants to isolate 32 strains bearing 19 mostly novel tubulin mutations that conferred differing degrees of resistance to two anti-tubulin compounds. The majority of the tubulin mutations were located outside of the drug-binding sites in the three-dimensional tubulin structure, suggesting that structural changes underlie the drug resistance conferred by these mutations. Thus, single-tubulin-gene disruptants are an efficient means of generating tubulin mutants for the study of the structure–function relationship of tubulin and for the development of novel therapies based on anti-tubulin agents.

## INTRODUCTION

Microtubules are fundamental cytoskeletal filaments that play pivotal roles in eukaryotic cell functions such as cell division, intra-cellular transport, cell shape development, and cilia and flagella assembly. Microtubules are produced by polymerization of α/β-tubulin heterodimers. Most eukaryotic cells possess multiple genes encoding α- and β-tubulin. For example, humans possess seven genes that encode α-tubulin and eight genes that encode β-tubulin, with each gene encoding a slightly different amino acid sequence. The presence of multiple genes for the two types of tubulin makes it difficult to study the properties of a particular tubulin species by genetic analysis, because the effects arising from mutation of one of the genes can be masked by the expression of the remaining intact genes.

The single-cell green alga *Chlamydomonas reinhardtii* is a useful experimental organism for studying tubulin function because it possesses a small number of tubulin genes and it produces microtubule-based organelles, flagella. In addition, there is a wide range of genetic tools available and a large amount of biological data has been accumulated for this species. In contrast to the majority of eukaryotes, *C. reinhardtii* possesses only two genes (*tua1* and *tua2*) encoding α-tubulin and two genes (*tub1* and *tub2*) encoding β-tubulin (1, 2). The two genes for each type of tubulin encode the same amino acid sequence (2, 3), and the expression of all four genes is up-regulated after flagellar excision (4). Whether the two genes in each pair are expressed independently of each other has not yet been firmly established, but the genes do appear to be expressed indiscriminately during flagella formation (4).

Although *C. reinhardtii* possess only two genes for each tubulin, the presence of more than one gene expressing the same protein still makes it difficult to isolate tubulin mutants. To date, only five kinds of tubulin mutations have been reported: a *tua1* mutation (Y24H) that confers amiprophos-methyl (APM) and oryzalin resistance (upA12) (3); two kinds of mutations in *tua2* (D205N and A208T) that confer colchicine hypersensitivity (*tua2-1* etc., suppressors of *uni-3-1*, a mutant lacking δ-tubulin) (5); and two mutations in *tub2* (K350E and K350M) that confer colchicine resistance (*col*^*R*^*4* and *col*^*R*^*15*) (6). APM, oryzalin, colchicine, and propyzamide are compounds that inhibit tubulin polymerization. These compounds other than colchicine inhibit plant tubulin polymerization at low concentrations and are used as herbicides.

Here, we isolated eight *tua2, tub1*, or *tub2* disruptants from an insertional library comprising around 8000 clones (7). We also obtained a *tua1* disruptant from the *Chlamydomonas* Library Project (CLiP) (8). We then used one of the *tub2* disruptants and two double-disruptants possessing only one α-tubulin gene and one β-tubulin gene as parent strains for the production of 32 mutants showing various degrees of resistance to propyzamide and oryzalin. Thus, the use of single-tubulin-gene *C. reinhardtii* disruptants enabled efficient isolation of a large number of tubulin mutants resistant to anti-tubulin agents.

## RESULTS

### Isolation of tubulin-gene disruptants

A library of around 8000 clones was constructed by inserting the *aphVIII* gene cassette into the genome of *C. reinhardtii* (7). Then, the library was screened by PCR using primer pairs consisting of one primer targeting a consensus sequence of the four tubulin genes and another primer targeting the *aphVIII* fragment. As a result, we isolated eight tubulin gene disruptants: three showing *tua2* disruption (*tua2-A, tua2-B, tua2-C*), two showing *tub1* disruption (*tub1-A, tub1-B*), and three showing *tub2* disruption (*tub2-A, tub2-B, tub2-C*). Fig. S1A shows the sites of the *aphVIII* cassette insertion in the eight disruptants, and Fig. S1B shows the PCR confirmation of the structure of the disrupted genes. In six of the disruptants (*tua2-A, tua2-B, tua2-C, tub1-B, tub2-A, tub2-C*), the *aphVIII* cassette was inserted into the gene. In the remaining two disruptants, the *aphVIII* cassette was inserted after the open reading frame (*tub1-A*) or within an intron (*tub2-B*). In all eight disruptants, *AphVIII* cassette insertion resulted in total disruption of mRNA expression of the affected tubulin gene, as confirmed by northern blot analysis (Fig. S2A). For *tua1-A, tua2-A, tub1-B*, and *tub2-A*, semi-quantitative real-time PCR was performed and again no expression of mRNA from the tubulin genes was detected (Fig. S2B).

None of the disruptants exhibited any apparent defects in growth rate (data not shown), tubulin expression (Fig. S2C), or flagellar regeneration after amputation (Fig. S2D), suggesting that the disruptants still produced sufficient α/β-tubulin heterodimer for their cellular functions via the remaining intact genes. The mean flagellar length was comparable among the disruptants (see Fig. S2D). The five β-tubulin disruptants showed some difference in their sensitivity to colchicine: *tub1-A* and *tub1-B* showed stronger resistance while *tub2-A* showed weaker resistance than wild type (Fig. 1), although the sensitivity somewhat varied among alleles (Fig. 1).

**Figure 1.**
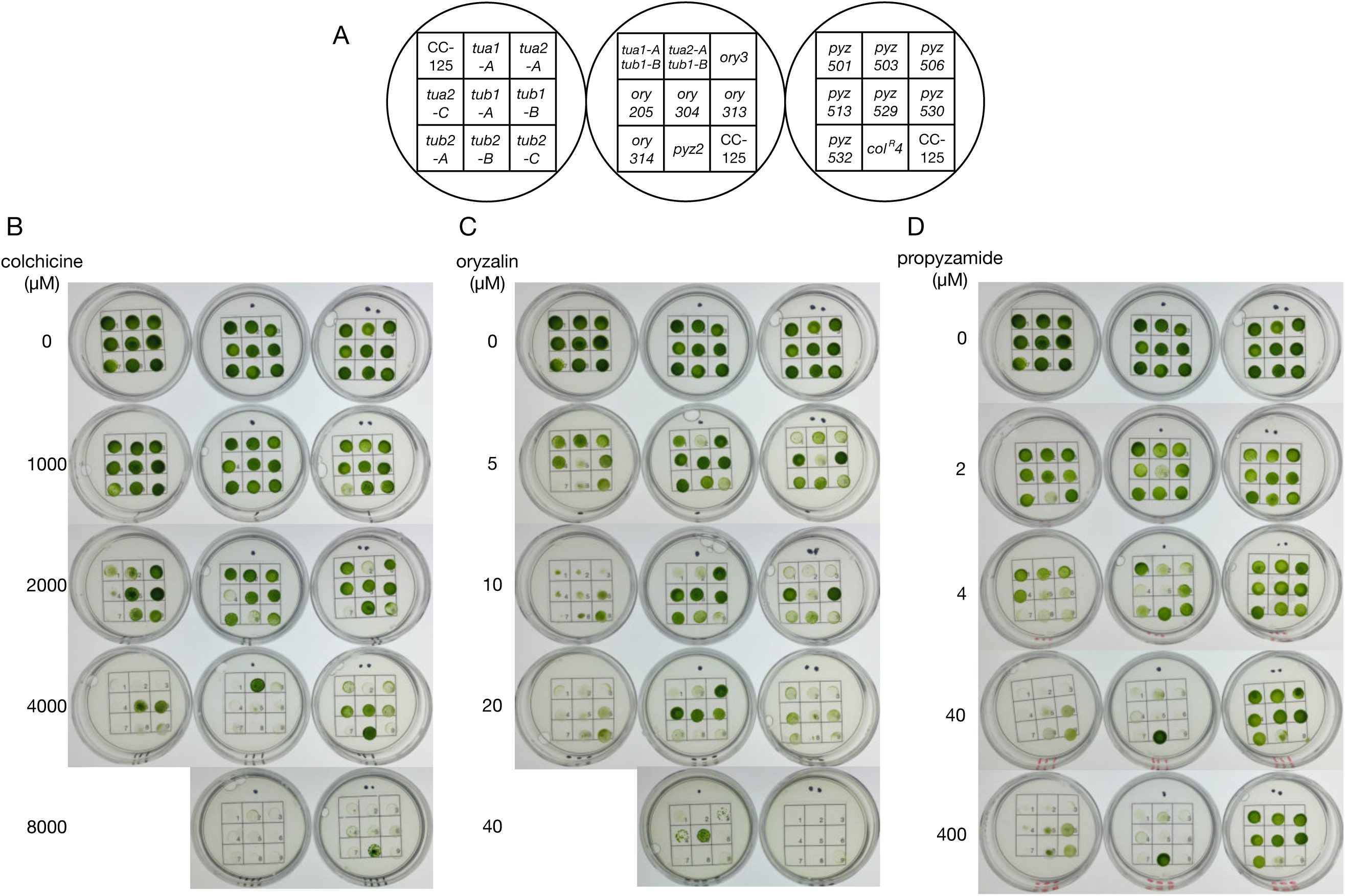
Drug sensitivities of tubulin-gene disruptants and missense mutant strains. Cell growth on TAP agar plates containing different concentrations of anti-tubulin drugs. Three plates each inoculated with nine strains were used for each condition, as summarized in panel (A). Sensitivity to colchicine (B), oryzalin (C), and propyzamide (D) was examined.

### Mutant isolation using tubulin-gene disruptants

Next, we used the disruptants to isolate *C. reinhardtii* strains expressing tubulins with missense mutations. Three parent strains were used: *tub2-A*, a double disruptant generated by crossing *tua1-A* with *tub1-B*, and a double disruptant generated by crossing *tua2-A* and *tub1-B*. As a result of 1-3 trials with each parental strain against oryzalin or propyzamide, 32 strains showing a total of 19 different tubulin missense mutations were isolated. Table 1 shows the obtained mutants classified by the gene affected, as well as the results of a qualitative assessment of each strain’s resistance to oryzalin and propyzamide. Most of the oryzalin-resistant strains, as well as a propyzamide-resistant mutant (*pyz532*), had a missense mutation in an α-tubulin gene. In contrast, most of the propyzamide-resistant strains, other than *pyz532*, had mutations in a β-tubulin gene.

**Table 1.** Tubulin missense mutants isolated in this study.

| Gene altered | Strain | Mutation (DNA) | Mutation (protein) | Resistance to |  |  |
| --- | --- | --- | --- | --- | --- | --- |
|  |  |  |  | oryzalin | propyzamide | colchicine |
| *tua1* | *ory2* | TAC->CAC | Y24H | +++ | +/- | +/- |
|  | *ory3* | TTC->CTC | F52L | ++++ | -- | + |
|  | *ory205, ory505* | TCC->GCC | S165A | ++++ | -- | -- |
|  | *pyz532* | TTC->TTA | F351L | + | +++ | --- |
| *tua2* | *ory304* | TAC->AAC | Y24N | ++++ | --- | +/- |
|  | *ory314* | TTC->TGC | F49C | +++ | - | +/- |
|  | *ory313* | TTC->GTC | F138V | ++++ | +/- | +/- |
| *tub1* | *pyz530, pyz534* | GAG->AAG | E198K | +++ | +++ | ++ |
|  | *pyz529, pyz531, pyz533* | TTC->TGC | F266C | --- | +++ | +++ |
| *tub2* | *pyz2, 524* | CAG->CAT | Q134H | ++ | +++ | - |
|  | *pyz506* | CTG->CAG | L165Q | +/- | +++ | + |
|  | *pyz502, pyz525, pyz526, pyz527* | GAG->AAG | E198K | N.E. | N.E. | N.E. |
|  | *pyz503* | GAG->TTG | E198L | +/- | +++ | -- |
|  | *pyz513* | ATC->AAT | I236N* | +++ | +++ | +++ |
|  | *pyz536* | TTC->TGC | F266C | N. E. | N.E. | N.E. |
|  | *pyz8, pyz9, pyz523* | AAG->GAG | K350E | + | + | ++++ |
|  | *pyz6* | AAG->ATG | K350M | N.E. | N.E. | N.E. |
|  | *pyz528* | AAG->AAT | K350N** | N.E. | N.E. | N.E. |
|  | *pyz501, pyz504, pyz514, pyz535* | ATC->TTC | I368F | --- | +++ | + |

N.E., not examined; +, resistant; -, sensitive; +/-, wild-type level. The number of + and - symbols represents the qualitative difference among the strains. \*, mutation found in human cancer cells resistant to 2-methoxyestradiol(23).\*\*, mutation found in human cancer cells resistant to indanocine and 2-methoxyestradiol(25, 33).

Figure 2 shows a predicted three-dimensional structure of *C. reinhardtii* α/β-tubulin heterodimer labeled with the site of each missense mutation reported here and in previous studies (3, 5, 6). Five of the isolates had mutations that have been reported previously: *ory2* had a *tua1* Y24H mutation as did upA12 (3); *pyz8, pyz9*, and *pyz523* had a *tub2* K350E mutation as did *col*^*R*^*4* (6); and *pyz6* had a *tub2* K350M mutation as did *col*^*R*^*15* (6). The five mutants isolated in the present study exhibited stronger drug-resistance than the three previously reported mutants (data not shown). This stronger drug-resistance may reflect the fact that the mutants isolated here express only mutated α- or β-tubulin from a single gene, whereas previously reported mutants express a mutated tubulin together with a wild-type counterpart.

**Figure 2.**
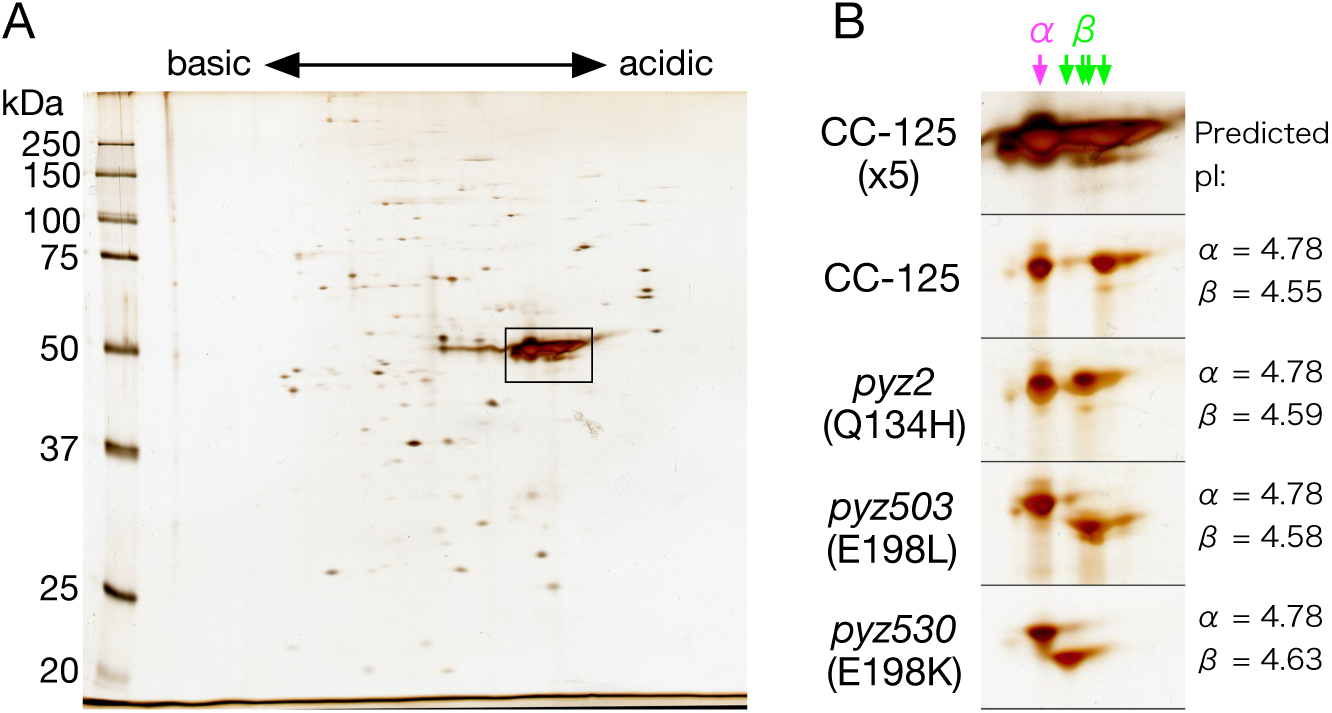
Two-dimensional polyacrylamide gel electrophoresis analysis of axonemes from strains *pyz2, pyz503*, and *pyz530*. Protein extracts of axonemes of wild type (CC-125), *pyz2, pyz503*, and *pyz530* were loaded on a two-dimensional polyacrylamide gel and stained with silver. pH range: 4.0–7.0. (A) Electrophoresis pattern of wild-type axoneme (∼10 μg loaded). (B) Portions of polyacrylamide gels showing the major spots of α- and β-tubulin. Upper panel shows a close-up of the area indicated by the box in (A). The lower four panels show the polyacrylamide gels after loading approximately 2 μg of axoneme. The predicted pIs of the wild-type and three mutant α- and β-tubulins are indicated to the right of the panels.

Some of the identified mutations involved the substitution of amino acids with different charges. For example, the propyzamide-resistant missense strains *pyz2/pyz524, pyz503*, and *pyz530/pyz534/pyz502/pyz525/pyz526/pyz527* expressed β-tubulins with the mutations Q134H, E198L, and E198K, respectively. The isoelectric point (pI) values of these β-tubulins predicted from their amino acid sequences were 4.59, 4.58, and 4.63, respectively, which were greater than the pI of wild-type β-tubulin (4.55). We confirmed the expression of β-tubulins with different pIs in those strains by two-dimensional polyacrylamide gel electrophoresis (2D-PAGE) of axonemal proteins from the mutants and wild type (Fig. 2). As expected, the spot of β-tubulin appeared at higher pH values in the order *pyz530* (E198K) > *pyz2* (Q134H) > *pyz503* (E198L) > wild type. The 2D-PAGE analysis also verified that each mutant expressed β-tubulin from only a single gene, since it detected no β-tubulin spots with the wild-type pI in mutant samples.

### Novel tubulin mutant strains exhibited various sensitivities to anti-tubulin agents

The mutant strains displayed various patterns of sensitivity to anti-tubulin agents (Table 1, Fig. 1). Several strains showed high oryzalin resistance in the order *ory304* > *ory3* > *ory205* > *ory313* > *ory314*. Three of these strains also showed hypersensitivity to propyzamide in the order *ory304* > *ory205* > *ory3*, and one of these strains, *ory205*, was also hypersensitive to colchicine. Several strains (*pyz2, pyz501, pyz503, pyz506, pyz513, pyz529, pyz530*, and *pyz532*) showed strong propyzamide resistance, remaining viable on a Tris–acetate–phosphate (TAP) agar plate containing more than 400 μM propyzamide whereas wild-type *C. reinhardtii* (CC-125) was barely viable at 40 μM. Of these eight mutant strains, three showed hypersensitivity to colchicine (*pyz532* > *pyz503* > *pyz2*) and the remaining five showed resistance to colchicine (*pyz529, pyz513, pyz530, pyz501*, and *pyz506*). The different sensitivities to colchicine and propyzamide in these mutants are interesting because the two agents bind to almost the same position on the tubulin heterodimer (9). Four of the eight propyzamide-resistant mutants exhibited oryzalin resistance in the order *pyz530* > *pyz513* > *pyz2* > *pyz532*, and two, *pyz501* and *pyz529*, were hypersensitive to oryzalin.

## DISCUSSION

By screening an *AphVIII* insertional library, we isolated three disruptants lacking *tua2*, two lacking *tub1*, and three lacking *tub2*. All were most likely null mutants (Fig. S2A). Although these disruptants lacked one of their tubulin-encoding genes, their cytoplasmic tubulin levels remained normal (Fig. S2C), suggesting the presence of an auto-regulatory mechanism that maintains the tubulin mRNA level, as observed in other eukaryotic cells (10). Indeed, in *tua1-A, tub1-B*, and *tub2-A*, the mRNA expression level of the remaining α- or β-tubulin gene was increased approximately 2-fold compared with wild type (Fig. S2B). Also, flagellar length, ability to produce flagella after amputation (Fig. S2D), and overall cell growth rate did not noticeably differ from the wild-type growth rate (data not shown). Thus, although *C. reinhardtii* possesses two α-tubulin genes and two β-tubulin genes, a single gene for each type is enough to supply the tubulin necessary for its cellular functions. However, it should be noted that the present findings do not mean that the two genes for each tubulin have exactly the same function; rather, the two genes may differ from each other in a subtle manner. For example, we observed that whereas the *tub1* disruptants were resistant to colchicine, the *tub2* disruptants were sensitive although some allele-specific variation was observed (Fig. 1). This suggests that there is some difference in the regulation of gene expression that is dependent on the concentration of free tubulin in the cytoplasm (11). Thus, how the two genes encoding the two tubulins differ in their function and regulation warrants further investigation, and our single-tubulin-gene mutants established here should be useful for such investigations.

Next, we used the disruptants to obtain mutants with resistance to two anti-tubulin agents, propyzamide and oryzalin. Several rounds of trials to isolate mutants resistant to one or both of the agents afforded 8 mutants with 7 different α-tubulin gene missense mutations and 24 mutants with 12 different β-tubulin gene missense mutations. The number of mutations obtained was much larger than the total number that has been reported previously (i.e., 3 kinds of α-tubulin mutations and 2 kinds of β-tubulin mutations)(3, 5, 6). In addition, we found that about one-third of the colonies picked from the screening plates harbored a mutation in a tubulin gene (data not shown). Together, these findings suggest that our approach of using single-tubulin-gene disruptants is a highly efficient means of obtaining tubulin mutant strains.

How the sensitivity to anti-tubulin agents varied in the mutants is an important issue that warrants clarification. Some of the tubulin mutants that conferred resistance to the anti-tubulin agents had a mutation near to where the anti-tubulin agents bind to the α/β-tubulin heterodimer (Fig. 3). The binding site of oryzalin, inferred from that of an analogous compound, tubulysin M, is at the intra-dimer interface (12) close to the α-tubulin mutation F351L (in strain *pyz532*). Other *ory* mutants whose mutation sites occur independently of the tubulysin M-binding site may confer oryzalin resistance by modulating the three-dimensional structure of α-tubulin. Likewise, a propyzamide-like compound, 2RR, is known to bind at the inter-dimer interface (13) close to several of the identified β-tubulin mutation sites: E198K/L (in *pyz530/pyz534/pyz502/pyz525/pyz526/pyz527* and *pyz503*), I236N (in *pyz513*), K350N/E/M (in *pyz528, col*^*R*^*4/ pyz8/pyz9/pyz523*, and *col*^*R*^*15/pyz6*), and I368F (in *pyz501/pyz504/pyz514/pyz535*). Other mutations may confer propyzamide resistance through some structural change in α/β-tubulin. Another interesting observation is that the β-tubulin mutation E198K conferred colchicine resistance whereas the E198L mutation conferred high colchicine sensitivity. This suggests that the electric charge of E198 is a critical determinant of colchicine sensitivity.

**Figure 3.**
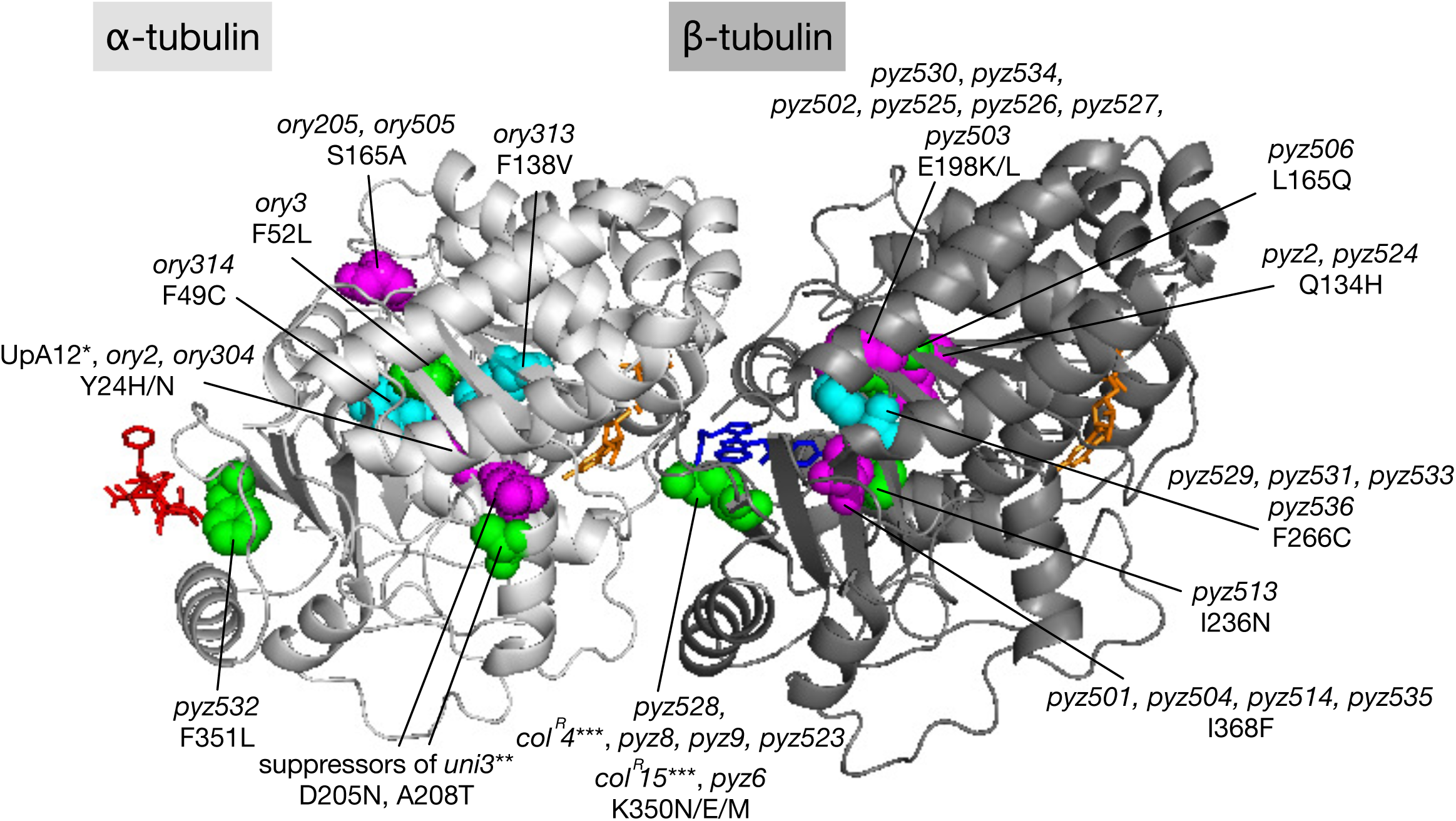
Predicted three-dimensional structure of *Chlamydomonas reinhardtii* α/β-tubulin heterodimer showing the mutations reported in the present and previous studies. Light gray, α-tubulin; dark gray, β-tubulin. Altered amino acids are shown as sphere representations. The binding sites of tubulysin M (red, an oryzalin-like compound) and 2RR (blue, 3-[(4-{1-[2-(4-aminophenyl)-2-oxoethyl]-1H-benzimidazol-2-yl}-1,2,5-oxadiazol-3-yl)a mino]propanenitrile, a propyzamide-like compound) were determined by applying the alignment command in MacPyMol software to the tubulin structures 4ZOL and 4O2A reported in the presence of these compounds (12, 13). The orange stick representations show GTP (in α-tubulin) and GDP (in β-tubulin). Mutations identical to previously reported mutations are marked with asterisks: *, (3); **, (5); and ***, (6).

Several of the mutations detected in the present study are similar to those reported in other organisms (Table S1). For α-tubulin, mutation F49C in *ory314*, F52L in *ory3*, and S165A in *ory205*/*ory505* have been reported in a *Toxoplasma gondii* oryzalin-resistant mutants (14, 15). For β-tubulin, mutation Q134H in *pyz2/pyz524* has been reported in a *Beauveria bassiana* benzimidazole-resistant mutant (16); mutations E198K/L in *pyz530/pyz534/pyz502/pyz525/pyz526/pyz527* and *pyz503* are found in fungi and nematodes that confer benzimidazole resistance and phenylcarbamate hypersensitivities (17-22); I236N in *pyz513* corresponded to the mutation responsible for resistance to the anti-cancer drug 2-methoxyestradiol in human epithelial cancer cells (23); K350N in *pyz528* corresponded to the mutations responsible for colcemid and vinblastine resistance in Chinese hamster ovary (CHO) cells (24) and indanocine resistance in human leukemia cells (25). Mutations Y24N, F138V, and F351L in α-tubulin (*ory304, ory313*, and *pyz532*) and mutations L165Q, F266C and I368F in β-tubulin (*pyz506, pyz529/pyz531/pyz533/pyz536*, and *pyz501/pyz504/pyz514/pyz535*) are being reported here for the first time; further investigations are needed to examine whether these mutations are responsible for altered drug sensitivity in other organisms.

Although the present study selected mutants based only on their resistance to two anti-tubulin agents, use of other agents such as the microtubule-stabilizing agent paclitaxel, or screening for other properties such as hypersensitivity to drugs, resistance to low temperature, or deficiency in flagellar formation and motility will lead to the isolation of a greater variety of mutants. Detailed analyses of many such mutants will deepen our understanding of the structure–function relation of tubulins. Since some of the tubulin mutations identified in the present study corresponded to mutations found in human tubulins that confer drug resistance in cancer cells, we expect that studies of *Chlamydomonas* tubulin mutants will contribute to the development of improved cancer therapies.

## MATERIALS AND METHODS

### Isolation of tubulin-gene disruptants

Eight tubulin-gene disruptants were isolated from a library of mutants generated by inserting the *aphVIII* gene (paromomycin resistance gene) into the genome of *C. reinhardtii* (7). Disruptants *tua2-B, tua2-C, tub1-B, tub2-A, tub2-B*, and *tub2-C* were isolated by performing PCR on the pooled transformants using primers targeting the *aphVIII* sequence (PSI103-F2 and RB02) and two tubulin consensus sequences (3’-Tus1891g and 3’-Tus1803g). A disruptant, *tub1-A*, was isolated using two alternative tubulin consensus primers (5’-Tus1082c and 5’-Tus1596g). A disruptant, *tua2-A*, was isolated using RB02 and an alternative consensus primer (3’-TuA2-3254g). Supplementary Information 1 shows the primers used in the present study. After screening, the disruptants were sequenced in the vicinity of their disrupted tubulin gene (Macrogen Japan Co., Japan).

In addition, a *tua1* disruptant (LMJ.RY0402.158052; referred to as *tua1-A* in the present study) was obtained from the *Chlamydomonas* Library Project (8); this disruptant has a long insertion composed of two facing paromomycin-resistant CIB1cassettes immediately before the stop codon in *tua1*.

The disruptants were backcrossed with wild-type *C. reinhardtii* (CC-125) and selected for tubulin-gene disruption by PCR before use. Double disruptants were constructed by standard methods (26), and selected from tetrads by PCR analysis.

### Isolation of anti-tubulin drug resistant mutants

*C. reinhardtii* strains whose *tub1* or *tub2* was disrupted with or without *tua2* disruption were grown to the mid-log phase and then irradiated by ultraviolet light until about 50% of the cells were killed. The culture was spread on TAP-agar plates containing 20 μM propyzamide or 10 μM oryzalin, kept in the dark for 12 h, and then incubated under light for 5–10 days. Colonies that appeared were transferred to liquid TAP medium in 96-well plates containing the same concentration of propyzamide or oryzalin. From each culture that grew, genomic DNA was extracted and subjected to PCR using the following primers: 5’-ChlaTuA1_long969 and 3’-TuA6260 (for *tua1*), 5’-Tua2-10g and 3’-TuA2-3288g (for *tua2*), 5’-tub1-33c and 3’-tub1-1667c (for *tub1*), and 5’-EcoTuB2-upper and 3’-XhoTuB2-lower (for *tub2*). The PCR products were processed for DNA sequencing (Macrogen Japan Co.).

### Drug-resistance test

*C. reinhardtii* strains were grown in liquid TAP medium until the mid-log phase, and then diluted or concentrated to 5 × 10^6^ cells/mL. Then, 3 μL of culture was spotted on a TAP-agar plate containing Propyzamide Reference Material (0–400 µM), Oryzalin Standard (0–40 µM), or colchicine (0–8000 µM) (all from Fujifilm Wako Pure Chemical Co., Japan) and cultured for a week at 26°C under 12-h light/12-h dark conditions. A wild-type strain (CC-125) and a colchicine-resistant mutant strain (*col*^*R*^*4* (6)) were used as references.

### Three-dimensional structure prediction of *C. reinhardtii* α/β-tubulin heterodimer

The three-dimensional structure of the *C. reinhardtii* α/β-tubulin heterodimer was predicted by using FAMS software (27) based on a known tubulin tetramer structure obtained from the Protein Data Bank (PDB ID: 1Z2B (28)). To determine the amino acids that most likely interacted with the examined drugs, *in silico* molecular docking analyses were performed using the ChooseLD program (29).

### 2D-PAGE of isolated axonemes

Axonemes were isolated from the *C. reinhardtii* strains by using standard procedures (30). A small aliquot of axonemal precipitate (∼2 or 10 μg) was extracted with a buffer containing 5 M urea and 2 M thiourea and analyzed by 2D-PAGE as described previously (31). Since α- and β-tubulin are modified post-translationally, the loading amount was adjusted so that their major forms only were detectable by silver staining. The predicted pI values of the wild-type and mutant tubulins were calculated by using the EMBOSS database and the Sequence Manipulation Suite, which is a collection of JavaScript programs for examining short protein sequences (https://www.bioinformatics.org/sms2/protein_iep.html) (32).

## ACKNOWLEDGMENTS

We thank Ms. Ai Tashiro, Haruka Fujisawa, and Tomoko Kurihara (Chuo University, Tokyo, Japan) for conducting the tubulin disruptant screening, and Ryosuke Suzuki, Hinano Sugai, Mami Ishigaki, and Yidong Tang (Chuo University) for performing the gene analysis of the tubulin mutants. We also thank Dr. Mitsuo Iwadate (Chuo University) for conducting the three-dimensional structure prediction and in silico molecular docking analyses, and Dr. Hiroko Kawai-Toyooka (The Univ. of Tokyo, Tokyo, Japan) for her technical support with the real-time polymerase chain reaction analyses.

## AUTHOR CONTRIBUTIONS

TKM and RK designed and conducted the research and wrote the paper. YO performed the real-time polymerase chain reaction analysis. TY and HF produced a temporal library of *Chlamydomonas reinhardtii* carrying the *aphVIII* gene.

